# Isolation and characterization of human monoclonal antibodies to pneumococcal capsular polysaccharide 3

**DOI:** 10.1101/2021.08.02.454853

**Authors:** Rachelle Babb, Christopher R Doyle, Liise-anne Pirofski

## Abstract

The current pneumococcal capsular polysaccharide (PPS) conjugate vaccine (PCV13) is less effective against *Streptococcus pneumoniae* serotype 3 (ST3), which remains a major cause of pneumococcal disease and mortality. Therefore, dissecting structure-function relationships of human PPS3 antibodies may reveal characteristics of protective antibodies. Using flow cytometry, we isolated PPS3-binding memory B cells from pneumococcal vaccine recipients and generated seven human PPS3-specific monoclonal antibodies (humAbs). Five humAbs displayed ST3 opsonophagocytic activity, four induced ST3 agglutination *in vit*ro, and four mediated both activities. For two humAbs, C10 and C27, that used the same variable heavy (V_H_) and light (V_L_) chain domains (V_H_3-9*01/V_L_2-14*03), C10 had fewer V_L_ somatic mutations, higher PPS3 affinity, more ST3 opsonophagocytic and agglutinating activity, whilst both humAbs altered ST3 gene expression *in vitro*. After V_L_ swaps, C10V_H_/C27V_L_ exhibited reduced ST3 binding and agglutination, but C27V_H_/C10V_L_ binding was unchanged. In C57Bl/6 mice, C10 and C27 reduced nasopharyngeal colonization with ST3 A66 and a clinical strain, B2, and prolonged survival following lethal A66 intraperitoneal infection, but only C10 protected against lethal intranasal infection with the clinical strain. Our findings, associate efficacy of PPS3-specific humAbs with ST3 agglutination and opsonophagocytic activity and reveal an unexpected role for the V_L_ in functional activity *in vitro* and *in vivo*. These findings also provide insights that may inform antibody-based therapy and identification of surrogates of vaccine efficacy against ST3.

**IMPORTANCE:** Despite the global success of pneumococcal conjugate vaccination, serotype 3 (ST3) pneumococcus remains a leading cause of morbidity and mortality. In comparison to other vaccine-included serotypes, the ST3 pneumococcal capsular polysaccharide (PPS3) induces a weaker opsonophagocytic response, which is considered a correlate of vaccine efficacy. Previous studies of mouse PPS3 monoclonal antibodies identified ST3 agglutination as a correlate of reduced ST3 nasopharyngeal colonization in mice, however neither the agglutinating ability of human vaccine-elicited PPS3 antibodies nor their ability to prevent experimental murine nasopharyngeal colonization has been studied. We generated and analysed the functional and *in vivo* efficacy of human vaccine-elicited PPS3 monoclonal antibodies and found that ST3 agglutination associated with antibody affinity, protection *in vivo*, and limited somatic mutations in the light chain variable region. These findings provide new insights that may inform the development of antibody-based therapies and next generation vaccines for ST3.

## INTRODUCTION

The current pneumococcal capsular polysaccharide conjugate vaccine, PCV13 is less effective against *S. pneumoniae* serotype 3 (ST3) than other vaccine-included serotypes (ST)’s. As a result, ST3 is a major cause of pneumonia and mortality in adults and children (1–5). Ample clinical data show that efficacy of pneumococcal conjugate vaccination reflects vaccine-mediated prevention of pneumococcal colonization and transmission, with vaccine-elicited ST-specific opsonophagocytic serum antibodies generally considered a surrogate for vaccine efficacy (6–8). However, a relationship between vaccine-elicited opsonophagocytic antibodies and protection against ST3 has not been established. In addition, compared to other vaccine-included STs, the capsular polysaccharide of ST3 (PPS3) is poorly immunogenic and induces a weaker opsonophagocytic antibody response (2). This reduced immunogenicity has been attributed to the thick ST3 capsule (9) as well as the limited ability of PPS3 antibodies to clear ST3 via opsonophagocytosis *in vivo* due to large amounts of ST3 capsule shedding (10). Nevertheless, opsonophagocytic PPS3 mouse and human monoclonal antibodies (mAbs) that are protective in ST3 sepsis and pneumonia models in mice have been generated (11–15). Notably, an opsonophagocytic mAb that protected against ST3 sepsis and pneumonia did not reduce ST3 colonization, whereas a non-opsonic mAb that agglutinated ST3, reduced colonization, protected against sepsis and pneumonia and also altered ST3 gene expression *in vitro* and *in vivo* (11, 13, 16).

Bacterial agglutination, including that of the pneumococcus, is a long-recognized correlate of PPS antibody efficacy in experimental models (17, 18). Whilst mouse and human PPS3 mAbs elicited by an experimental PPS3-TT conjugate revealed that ST3 opsonophagocytosis and agglutination were mutually exclusive functions (11, 13, 16, 19), serum derived antibodies to ST4 and ST23 exhibited both opsonophagocytic and agglutinating functions (20). Consistent with the latter, among a set of 5 PPS3 mouse mAbs generated in response to a PPS3-KLH conjugate, 4 exhibited both opsonophagocytic and agglutinating activity and only one mediated opsonophagocytosis (21). These findings suggest that the nature of PPS3 antibodies that mediate opsonophagocytosis and agglutination versus those that mediate one function and not the other may differ.

Reduced efficacy of PPS3-specific antibodies against ST3 disease has been attributed to impaired opsonophagocytic clearance, and it has been estimated that approximately 8 times more antibody is required to confer protection against ST3 based on the calculated correlate of protection for other pneumococcal STs (2, 10). Thus, deciphering the structural and functional characteristics of human vaccine elicited PPS3 antibodies may advance understanding of vaccine failure and facilitate development of antibody-based therapies and next generation vaccines. To gain insight into the nature of human PPS3-binding antibodies, we generated PPS3 human mAbs (humAbs) from human pneumococcal vaccine recipients and determined their molecular derivation, PPS3 binding, and function *in vitro* and *in vivo*.

## RESULTS

### PPS3 humAbs use gene segments from the VH3 family

Seven PPS3-binding humAbs (PPS3 humAbs) were generated and tested for PPS3 binding by ELISA (Figure 1). C38 had the strongest binding to PPS3 (EC_50_= 0.09 μg/ml), followed by C34 (EC_50_= 0.21 μg/ml), and C10 (EC_50_= 0.24 μg/ml). Binding to a ST3 clinical strain, B2 was also similar by whole cell ELISA and immunofluorescence (Figure S1 & S2).

**Figure 1:**
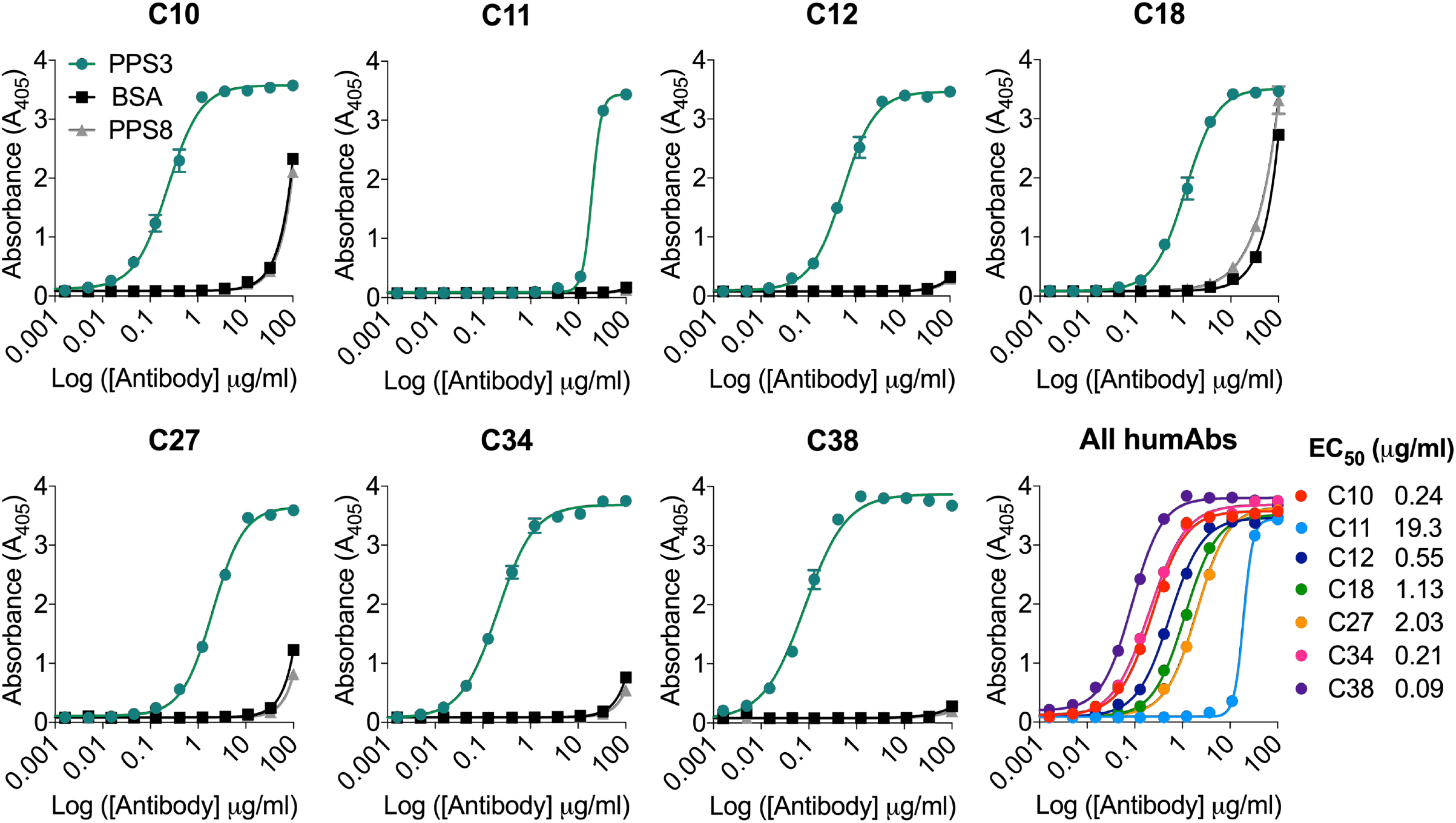
HumAb binding to pneumococcal polysaccharide 3 (PPS3) by ELISA. Binding as reflected by absorbance at 405 is shown on the Y axis for the humAb concentrations shown on the X axis for each humAb. Results are representative of 3 independent experiments (n = 2). The numerical half-maximal binding titer (EC_50_) for each humAb is indicated to the right of the panel depicting binding curves of all humAbs.

Sequencing analysis revealed that five humAbs (C10, C12, C27, C34, C38) used lambda light chains (LC)s and two (C11, C18) used kappa LCs. Based on IgBlast, six used variable heavy 3 (V_H_3) genes and one (C38) used a V_H_1 gene (Table 1). All seven humAbs had V_H_ and V_L_ CDR as well as FR somatic mutations (Figure S3 & S4). In addition, all seven humAb CDR3s differed by sequence and length, but four (C10, C27, C38, C11) had an Ala-Arg-Asp: ARD or Ala-Arg-Gly: ARG motif at the beginning of the V_H_ CDR3 region (Table 1). Two lambda humAbs, C10, C27 used the same heavy VDJ and LC VJ segments, but their FRs and CDRs differed by several somatic mutations (Figure 2). C10 and C27 had respectively, 9 and 8 V_H_ mutations conferring amino acid changes relative to germline IGHV3-9*01, including 4 at the same positions and a shared Lysine (K) in CDR2. C10 V_L_ was closer to germline IGVL2-14*03, with fewer mutations (5 versus 11) than C27, 4 of which were shared.

**Table 1.**
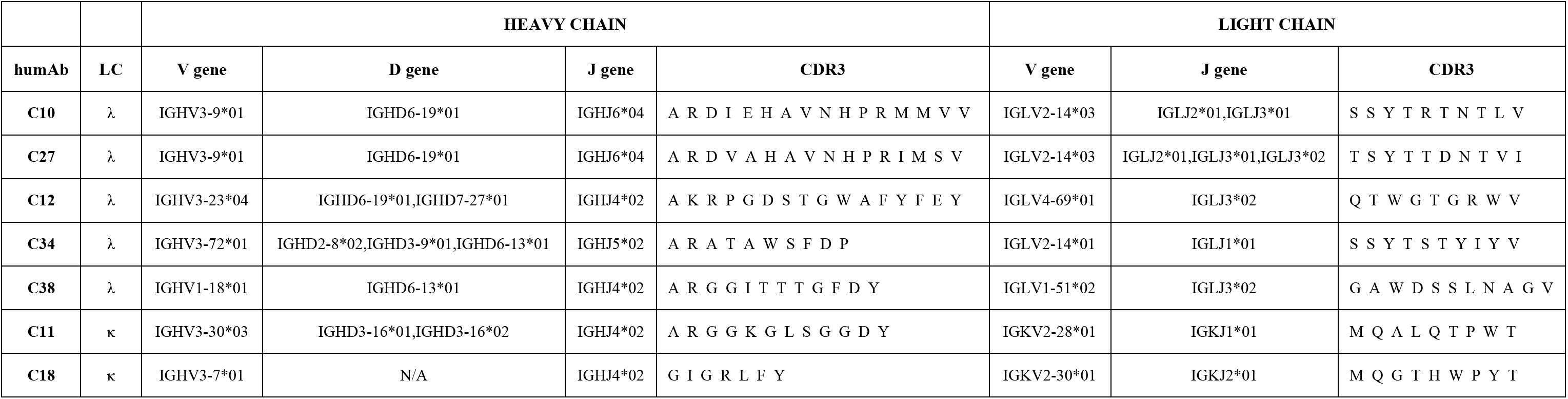
Heavy and light chain VDJ gene usage and CDR3 sequences for all PPS3 humAbs.

|  |  | HEAVY CHAIN |  |  |  | LIGHT CHAIN |  |  |
| --- | --- | --- | --- | --- | --- | --- | --- | --- |
| humAb | LC | V gene | D gene | J gene | CDR3 | V gene | J gene | CDR3 |
| C10 | λ | IGHV3-9*01 | IGHD6-19*01 | IGHJ6*04 | A R D I E H A V N H P R M M V V | IGLV2-14*03 | IGLJ2*01,IGLJ3*01 | S S Y T R T N T L V |
| C27 | λ | IGHV3-9*01 | IGHD6-19*01 | IGHJ6*04 | A R D V A H A V N H P R I M S V | IGLV2-14*03 | IGLJ2*01,IGLJ3*01,IGLJ3*02 | T S Y T T D N T V I |
| C12 | λ | IGHV3-23*04 | IGHD6-19*01,IGHD7-27*01 | IGHJ4*02 | A K R P G D S T G W A F Y F E Y | IGLV4-69*01 | IGLJ3*02 | Q T W G T G R W V |
| C34 | λ | IGHV3-72*01 | IGHD2-8*02,IGHD3-9*01,IGHD6-13*01 | IGHJ5*02 | A R A T A W S F D P | IGLV2-14*01 | IGLJ1*01 | S S Y T S T Y I Y V |
| C38 | λ | IGHV1-18*01 | IGHD6-13*01 | IGHJ4*02 | A R G G I T T T G F D Y | IGLV1-51*02 | IGLJ3*02 | G A W D S S L N A G V |
| C11 | κ | IGHV3-30*03 | IGHD3-16*01,IGHD3-16*02 | IGHJ4*02 | A R G G K G L S G G D Y | IGKV2-28*01 | IGKJ1*01 | M Q A L Q T P W T |
| C18 | κ | IGHV3-7*01 | N/A | IGHJ4*02 | G I G R L F Y | IGKV2-30*01 | IGKJ2*01 | M Q G T H W P Y T |

**Figure 2:**
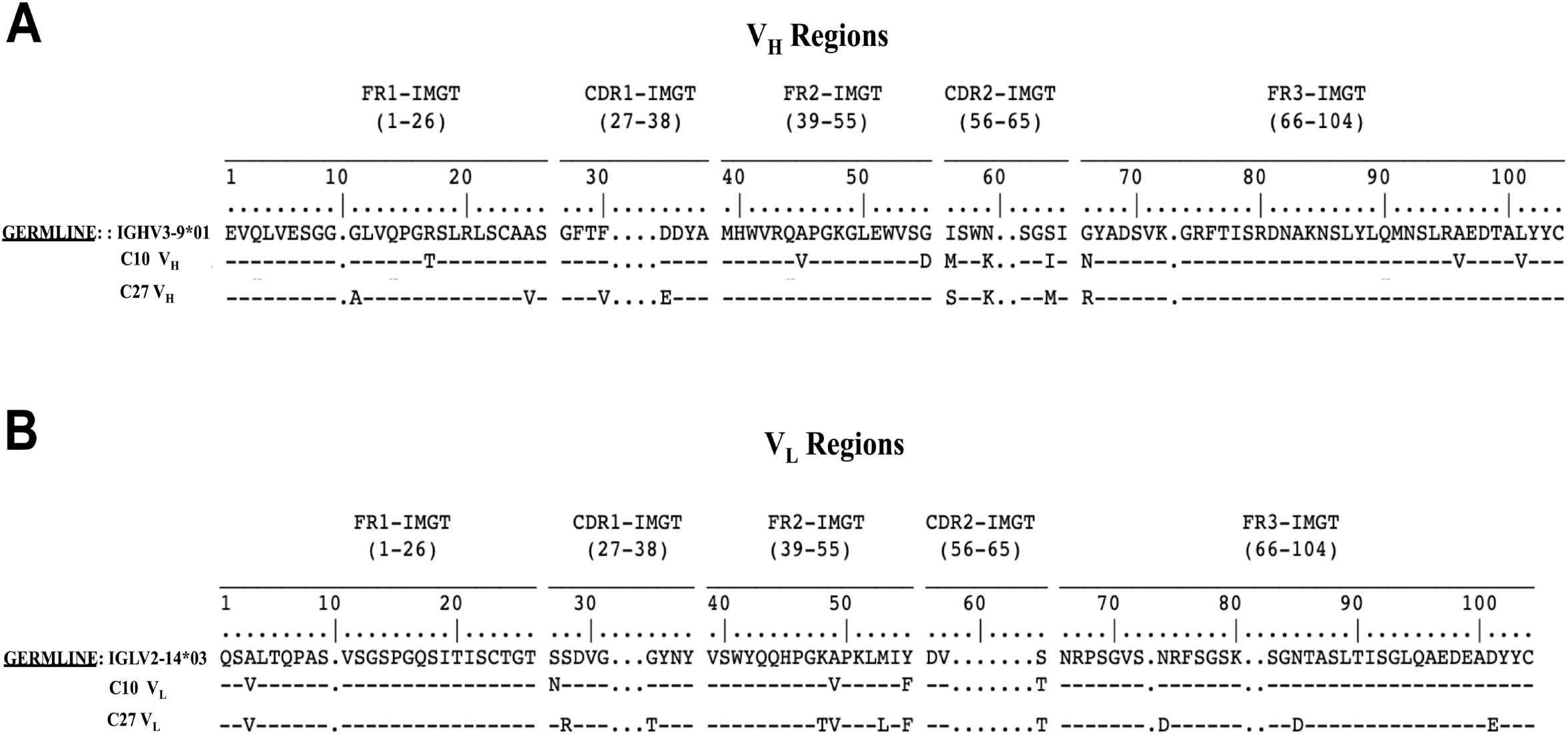
C10 and C27 variable heavy (V_H_) and light (V_L_) chain amino acid sequences. HumAb A) V_H_ and B) V_L_ sequences aligned with their germline counterparts based on IMGT/V-QUEST (sequence alignment software). Amino acid changes resulting from somatic mutations are indicated within the sequence alignment.

### PPS3 humAbs agglutinate ST3 *in vitro*

It has been previously reported that antibodies that agglutinate pneumococcus can reduce pneumococcal colonization (13, 22, 23). Thus, we determined the ability of the PPS3 humAbs to agglutinate ST3 A66 and the clinical strain, B2 by flow cytometry and validated our findings with light microscopy. C10, C12, C34 and C38 each exhibited dose-dependent agglutination of ST3. At 10 µg/ml, C34 and C38 agglutinated ∼75% and 89% of bacteria, respectively, whilst C10 and C12 agglutinated ∼48% and 39%, respectively (Figure 3A & B). Visual ST3 clumping was also observed with C10, C12, C34 and C38 by light microscopy (Figure 3C). Similar results regarding agglutination experiments were obtained with the clinical strain, B2 (Figure S5). F(ab’)_2_ fragments of C10 and C38 also agglutinated ST3 with levels comparable to their respective whole IgG (Figure 4A & B).

**Figure 3:**
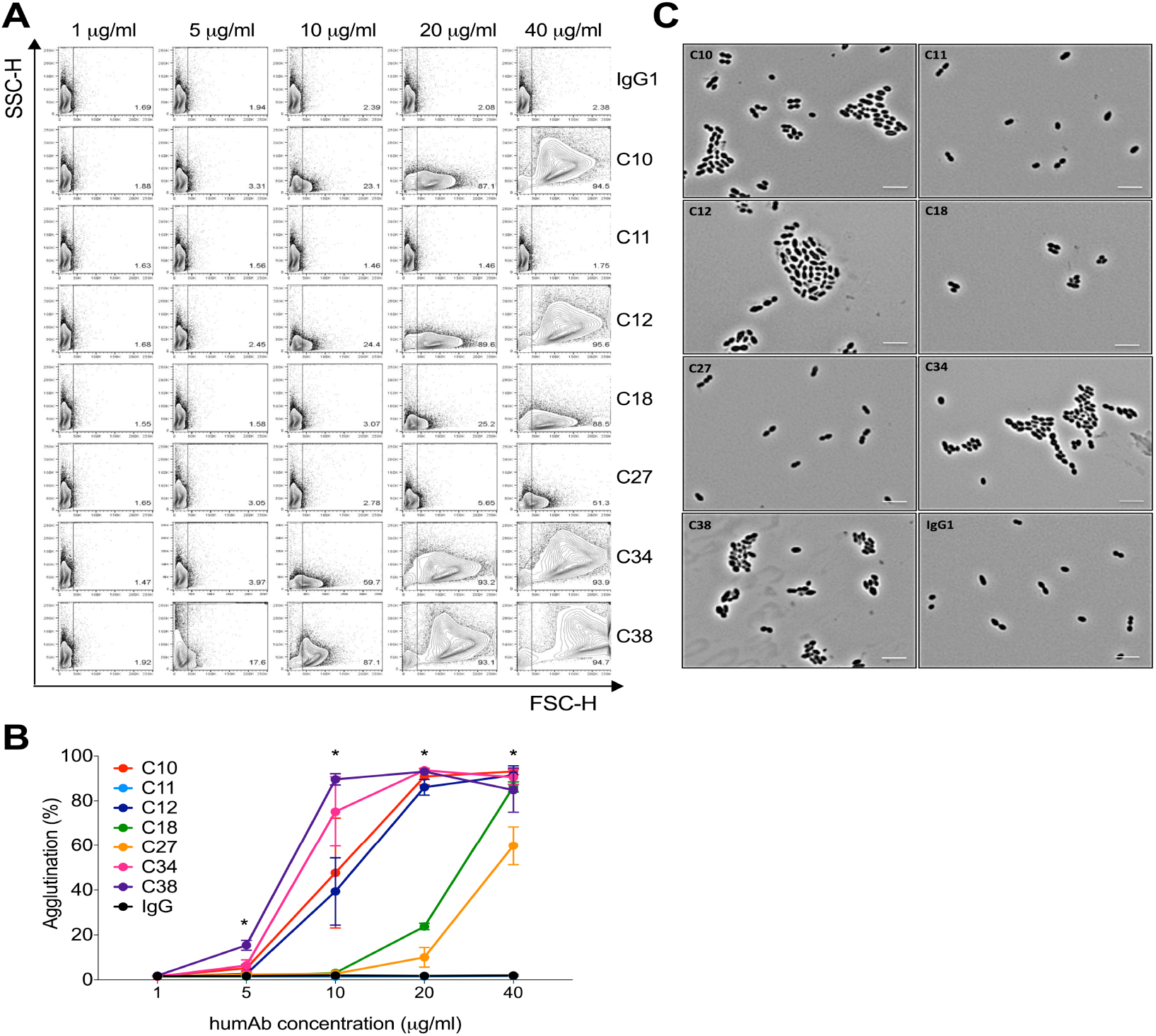
HumAb agglutination of ST3 A66. The ability of the humAbs to agglutinate ST3 (A66) was assessed by flow cytometry. A) Representative FACS dot plots showing the percentage agglutination of all humAbs and control human IgG1 at various concentrations by flow cytometry B) Percentage of agglutination is shown on the Y axis for different humAb concentrations indicated on the X axis. Graph represents data from 2 independent experiments (n = 2 per condition). C) Light microscopy images of humAbs (20 µg/ml) with ST3 A66. Images at 100x magnification are representative of 3 independent experiments (n = 2). Scale bars, 5 µm. By one-way ANOVA: At 5 µg/ml; (C38 vs IgG1 \*\*\**P*<0.001), at 10 µg/ml; (C34 & C38 vs IgG1 \**P*<0.05); at 20 µg/ml (C10, C12, C18, C34 & C38 vs IgG1 \*\**P*<0.01), at 40 µg/ml (C10, C12, C18, C27 & C38 vs IgG1 \*\*\**P*<0.001).

**Figure 4:**
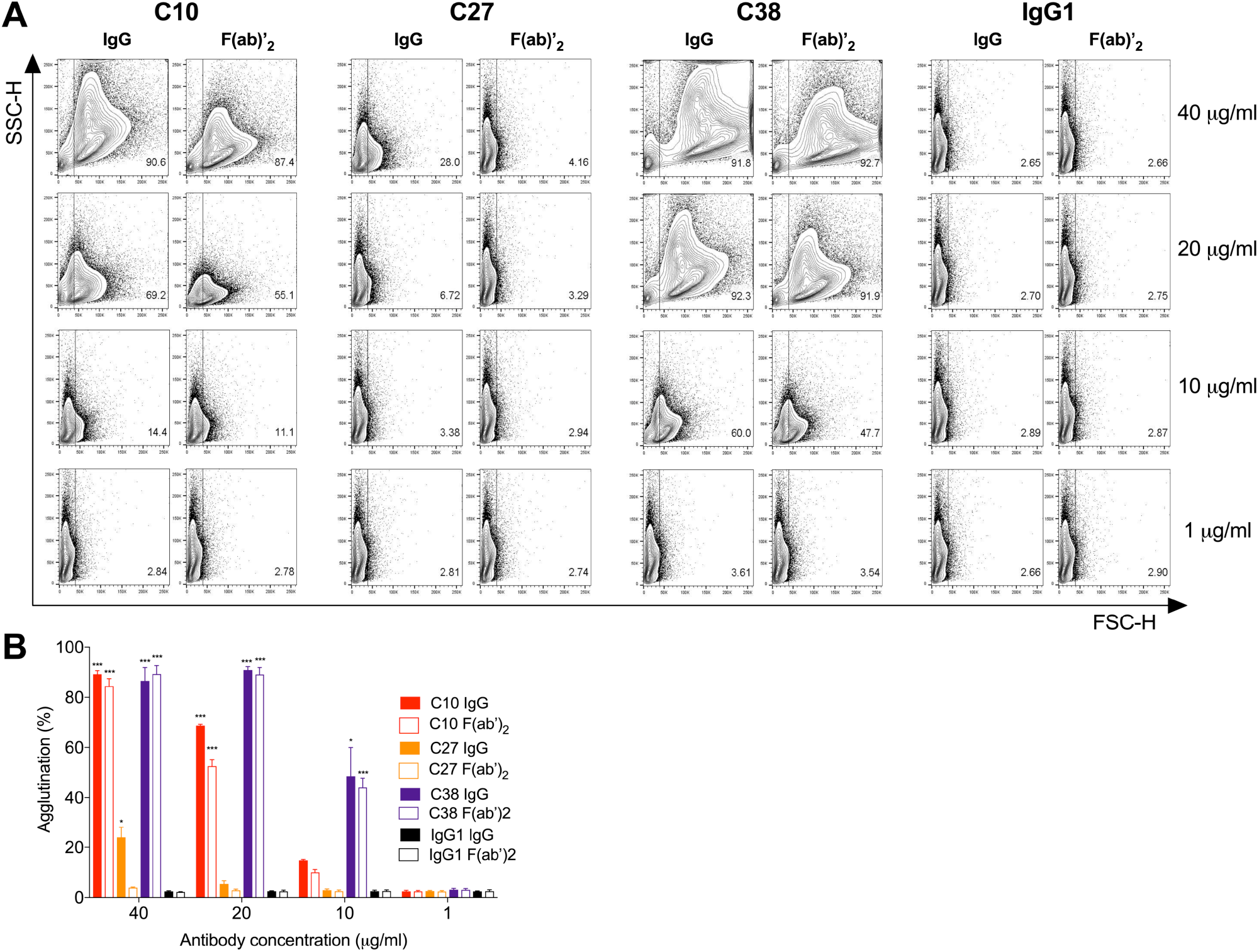
HumAb F(ab)’_2_ fragment agglutination of ST3 A66. The ability of whole IgG or F(ab’)_2_ fragments of humAbs (C10, C27, C38) to agglutinate ST3 (A66) was assessed by flow cytometry. A) Representative FACS dot plots showing the percentage agglutination of the indicated whole humAbs, F(ab)’_2_ fragments, or control IgG1 at various concentrations. B) Bar graph depicting percentage agglutination on the Y axis for whole humAb or F(ab’)_2_ fragment concentrations on the X axis. Results are representative of 2 independent experiments (n = 2 per condition). By one-way ANOVA: At 10 µg/ml; (C38 IgG, C38 F(ab’)_2_ vs their respective IgG1 controls \**P*<0.05); at 20 µg/ml; (C10 IgG, C10 F(ab’)_2_, C38 IgG, C38 F(ab’)_2_ vs their respective IgG1 controls \*\*\**P*<0.001); at 40 µg/ml; (C10 IgG, C10 F(ab’)_2_, C38 IgG, C38 F(ab’)_2_ vs their respective IgG1 controls \*\*\**P*<0.001).

### Opsonophagocytosis of ST3 by PPS3 humAbs

Functional activity of the humAbs was determined with the standard opsonophagocytic assay (OPA) used in the field (24, 25). C10 and C38 displayed the highest activity with significant reductions in CFU at 0.74 µg/ml (Figure 5) relative to the IgG1 control. C12, C18 and C34 reduced CFU at 2.2 µg/ml, and C11 and C27 at 20 µg/ml. When humAbs were incubated with ST3 without HL60 cells, C10, C18, C27, C34 and C38 reduced CFU relative to the control. This correlated with agglutination, except for C27.

**Figure 5:**
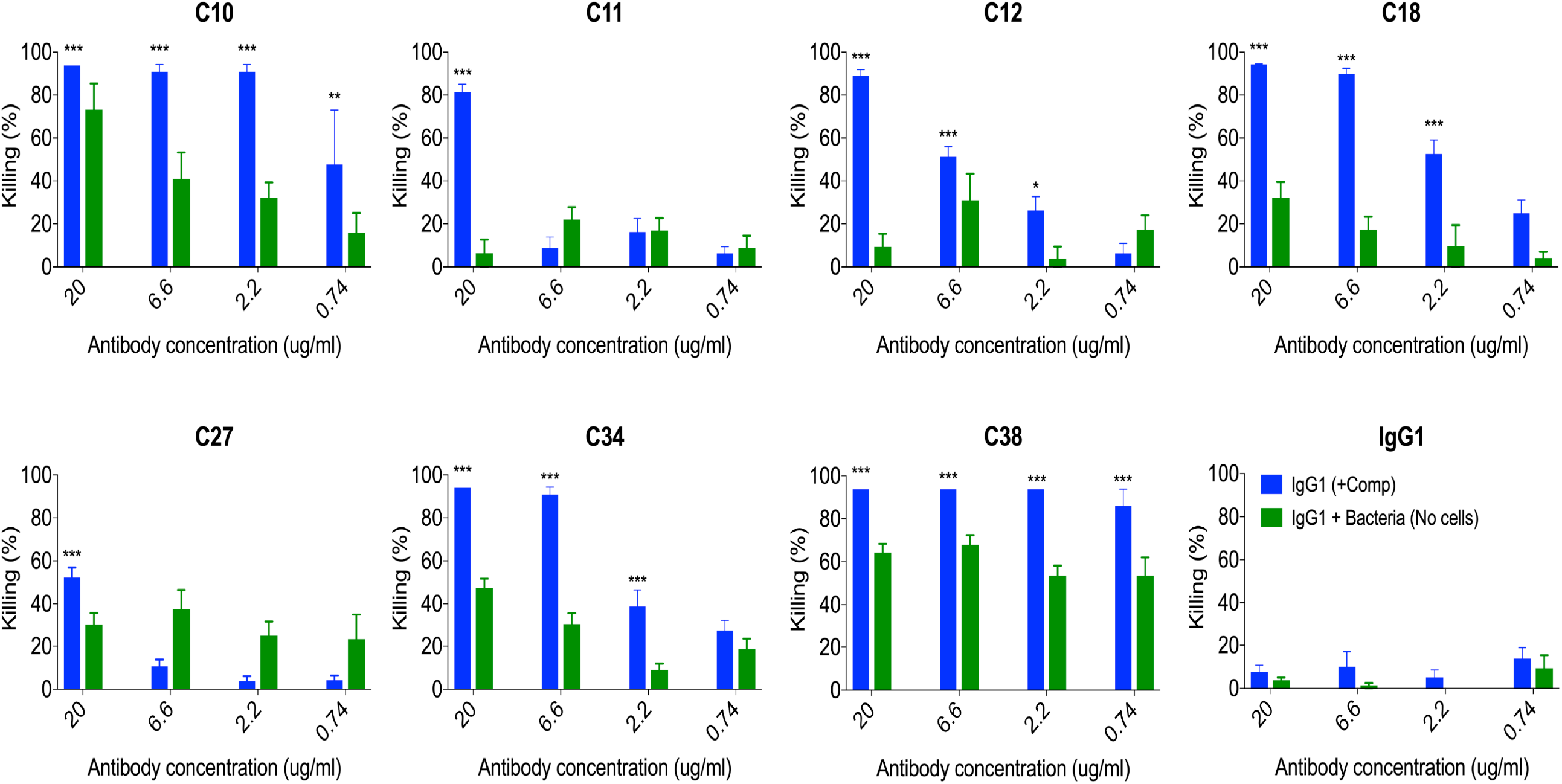
HumAb opsonophagocytic killing of ST3. HumAbs were tested for their opsonophagocytic killing activity with ST3 (A66) and HL60 cells. Percent killing is shown on the Y axis for the different humAb concentrations shown on the X axis. Results are representative of 2 independent experiments (n = 4 per condition). \**P*<0.05, \*\**P*<0.01, \*\*\**P*<0.001 (One-way ANOVA) for humAbs vs IgG1 control.

### PPS3 humAbs reduce A66 and B2 nasopharyngeal colonization in C57Bl/6 mice

We next performed nasopharyngeal (NP) colonization experiments in mice with C10 and C27. These humAbs were used because they use the same V_H_3-9*01/V_L_2-14*03 gene elements but have different affinities and functional activity *in vitro*. Compared to the IgG1 control, administration of C10 and C27 reduced NP CFU after infection with A66 (C10; P=0.0388, C27; P=0.0437) (Figure 6A) and B2 (C10; P=0.0128, C27; P=0.0015) (Figure 6B). CFU were not detected in the lungs (data not shown). Compared to IgG1-treated controls, B2-infected C10- and C27-treated mice had significantly lower TNF-α, IL-1α and IL-6 levels 4 days post infection (Figure 6C).

**Figure 6:**
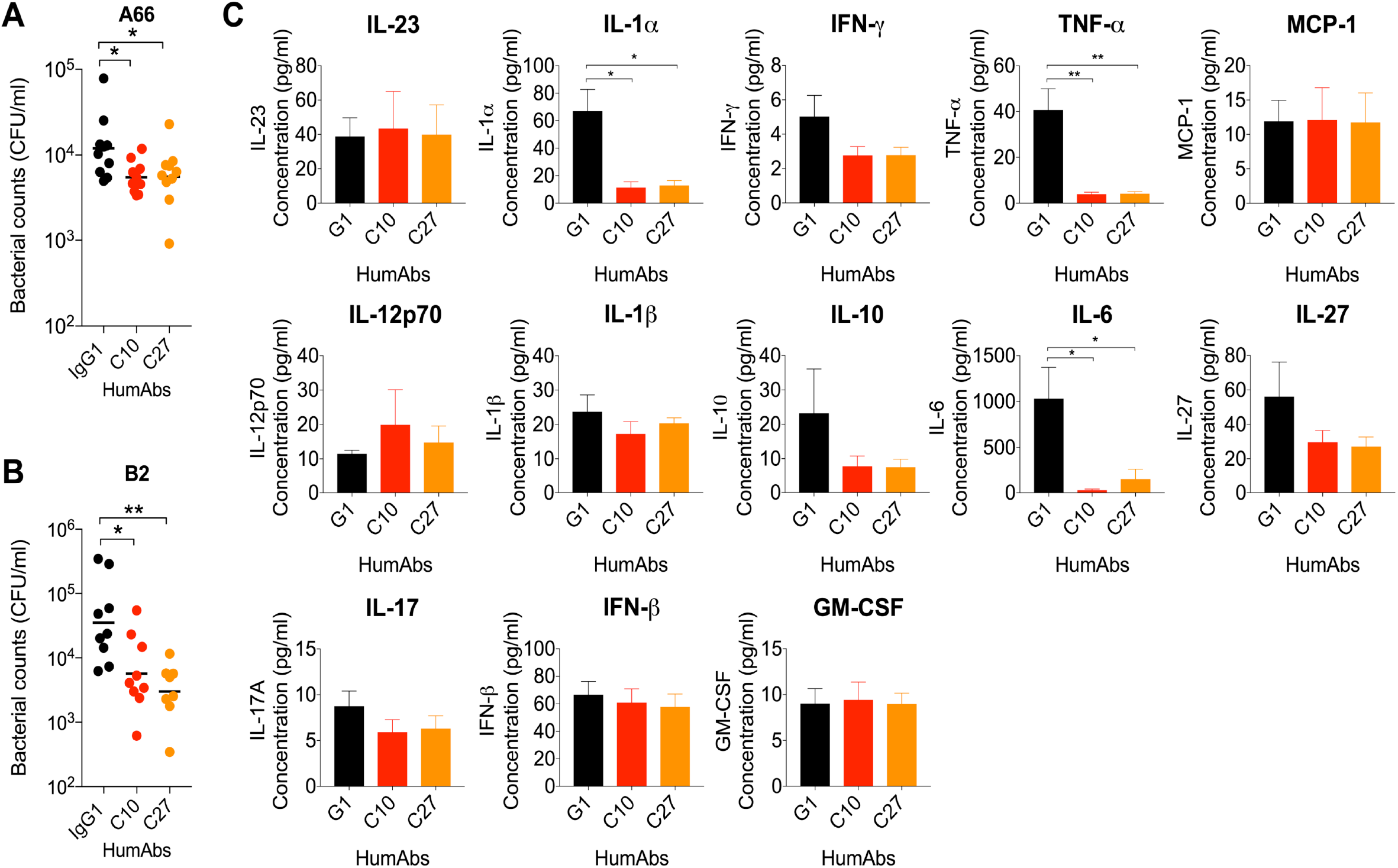
HumAb efficacy against ST3 colonization in C57Bl/6 mice. HumAbs or a control IgG1 were administered IN in C57Bl/6 mice 2 hrs before IN infection with A) 5 x 10^5^ CFU A66 or B) 1 x 10^7^ CFU B2. The nasal lavage CFU was enumerated 24 hours (A) or 4 days (B) post infection. CFU are depicted on the Y axis for humAbs shown on the X axis C) Indicated cytokine concentrations via legendplex 4 days after infection of C57Bl/6 mice with 1 x 10^7^ CFU B2 (B) are shown on the Y axis for the humAbs on the X axis. Results are representative of 2 independent experiments (n ≥ 5 mice/group). \**P*<0.05, \*\**P*<0.01, \*\*\**P*<0.001 (One-way ANOVA).

### PPS3 humAbs prolong survival of mice lethally infected with A66 and B2

The efficacy of C10, C27, and C38 was next investigated in lethal ST3 infection models. C38 was included because it exhibited strong ST3 binding, opsonophagocytosis, and agglutination. IP administration of all three mAbs prolonged survival after IP infection with A66 (Figure 7A). C10 was the most protective (92% survival, P=0.0001), followed by C27 (76%, P=0.001), and C38 (70%, P=0.0036). In a lethal IN infection model, IN administration of C10, but not C27 prolonged survival after infection with B2 (85%, P=0.0291) compared to the IgG1 control (Figure 7B).

**Figure 7:**
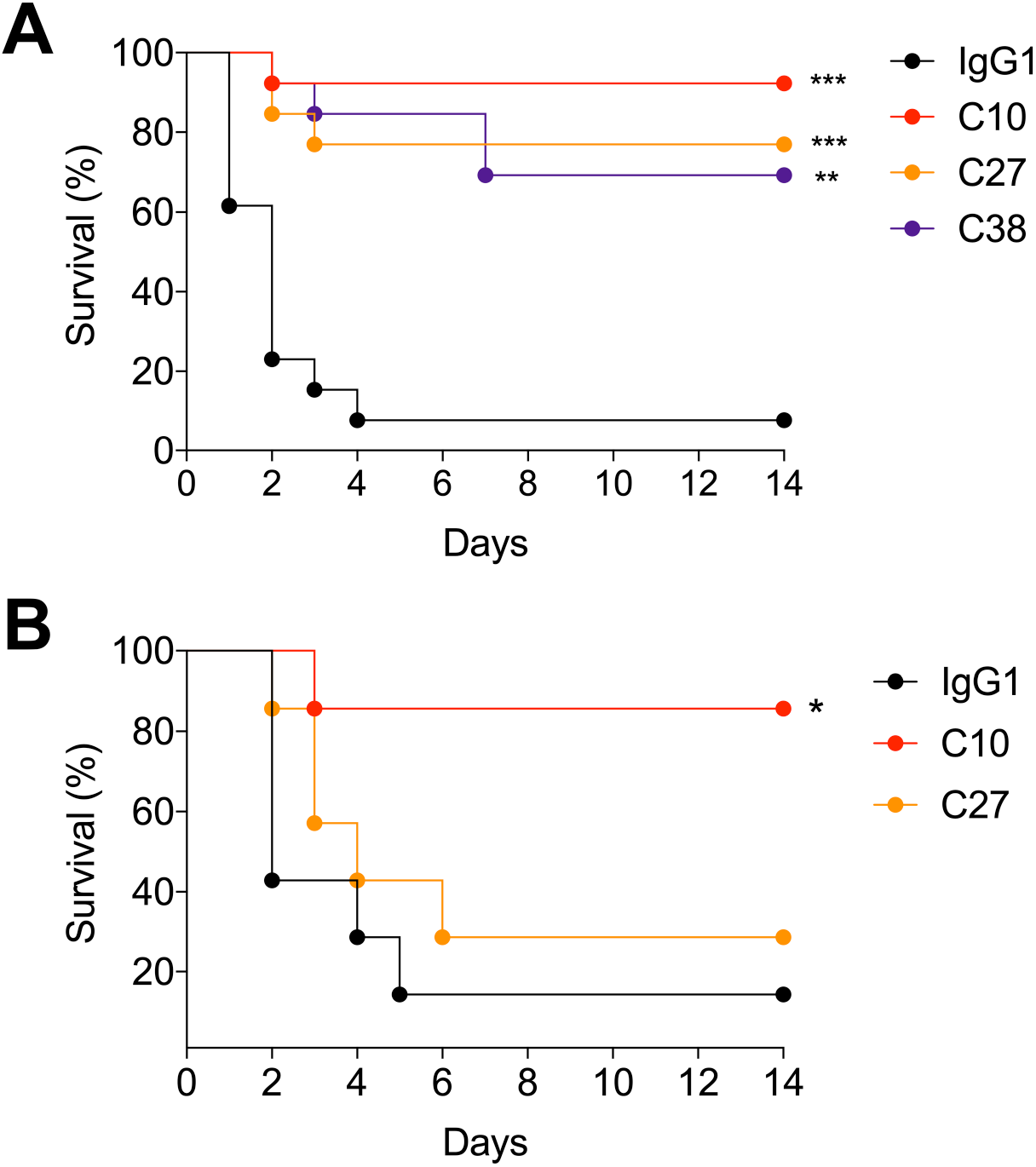
HumAb efficacy against lethal challenge with ST3 strains in C57Bl/6 mice. A) HumAbs or a control IgG1 were administered IP in C57Bl/6 mice 2 hrs before IP infection with 5x 10^5^ CFU A66 and then monitored for survival. B) HumAbs or a control IgG1 were administered IN in C57Bl/6 mice 2 hrs before IN infection with 5 x 10^7^ CFU B2 and monitored for survival. All curves show percent survival on the Y axis for the indicated humAbs monitored over 14 days shown on the X axis. Results are representative of 2 independent experiments (n ≥ 7 mice/group). \**P*<0.05, \*\**P*<0.01, \*\*\**P*<0.001, (Fisher’s exact test).

### HumAbs alter bacterial gene expression *in vitro*

Given that C27 did not promote agglutination or opsonophagocytosis *in vitro*, yet it reduced colonization and protected against lethal IP infection (sepsis), we sought an alternative mechanism by which it could mediate protection. Previous work showed that defined PPS3 mAbs enhanced ST3 A66 transformation frequency and competence, and one mAb, 1E2, altered ST3 gene expression *in vitro* and *in vivo* (13, 16, 19). Thus, we performed RT-qPCR on reactions of ST3 A66 incubated with C10 and C27 to analyze expression of ST3 genes that induce or respond to oxidative stress (*dpr*, *piuB*, *blpX*, *merR*, *comX*) and of which expression was altered in 1E2-treated mice following NP colonization (16). In comparison to an IgG1 control, C10 and C27 each induced a significant decrease in *dpr* gene expression (Figure 8). We also observed a decrease in *piuB*, *blpX*, *merR* and *comX* expression (Figure 8). There were no significant differences between C10 and C27 in the genes examined.

**Figure 8:**
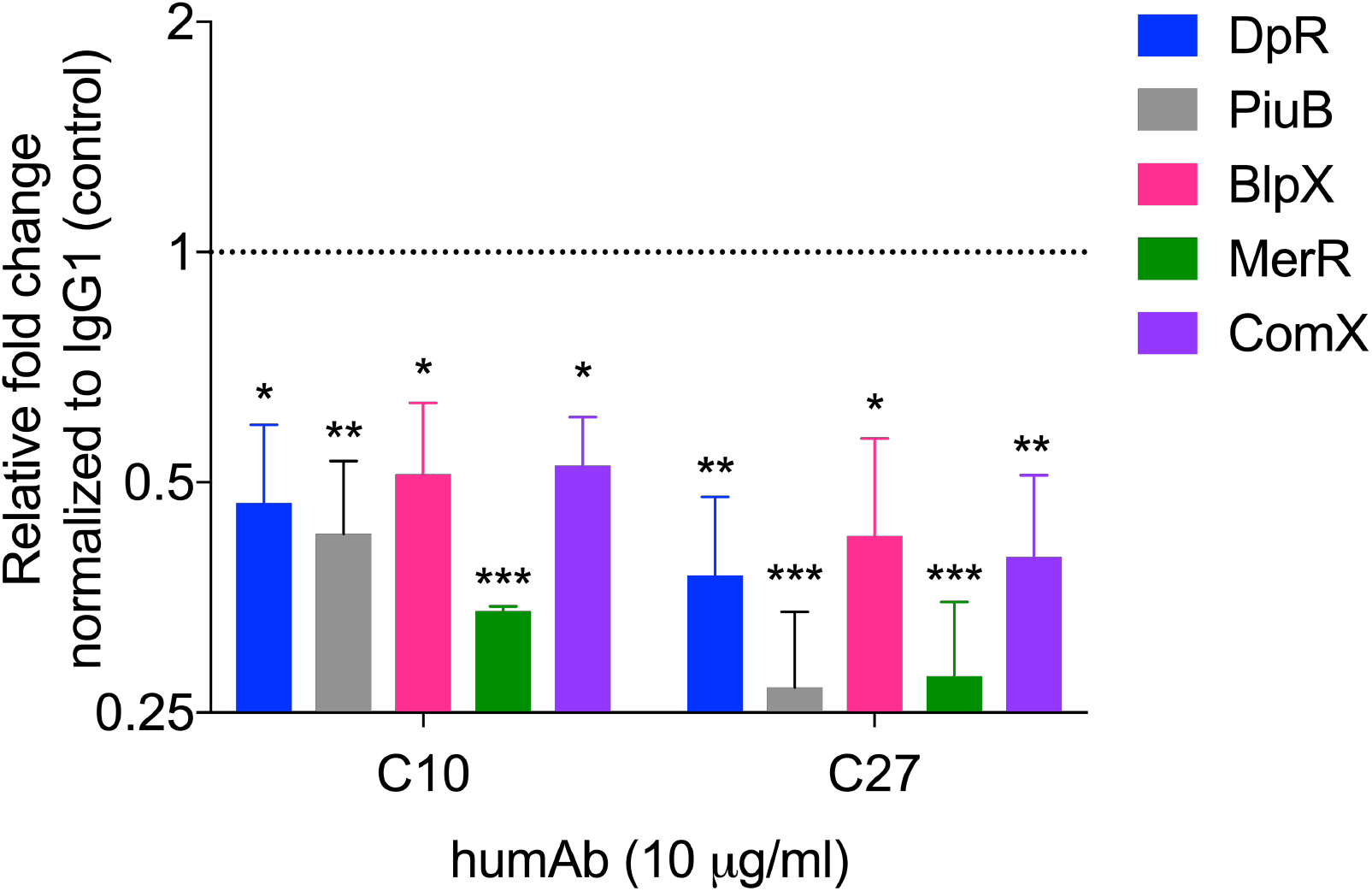
HumAbs mediate changes in expression of bacterial genes related to oxidative stress *in vitro*. The fold change in expression of the indicated genes in C10 or C27-treated bacteria relative to the control IgG1-treated bacteria was determined by RT-qPCR at 1.5 hours post-humAb addition. Relative expression of genes was determined using the Pfaffl method (55) (fold change is relative to the IgG1 control treated bacteria, expression =1). Data are pooled from 3 independent experiments, 3 samples per condition. \**P*<0.05, \*\**P*<0.01, \*\*\**P*<0.001, (One-way ANOVA) C10 or C27 vs IgG1.

### Analysis of humAbs with V_L_ swaps

Given that C10 and C27 use the same V_H_ and V_L_, but C27 had lower affinity, reduced ST3 binding and more mutations in its V_L_ region relative to the germline, we performed V_L_ swaps to evaluate the effect of V_L_ on binding and agglutination. PPS3 and B2 binding of C10 expressing the V_L_ of C27 (C10_H_C27_L_) was reduced compared to that of native C10, whereas C27 exhibited no differences in binding when expressing the C10 V_L_ (C27_H_C10_L_) (Figure 9A). In agglutination experiments with B2, 20 µg/ml of C10 promoted strong agglutination (∼75%) compared to C10_H_C27_L_ (∼10%), but there were no differences in agglutination for C27_H_C10_L_ relative to native C27 (Figure 9B & C).

**Figure 9:**
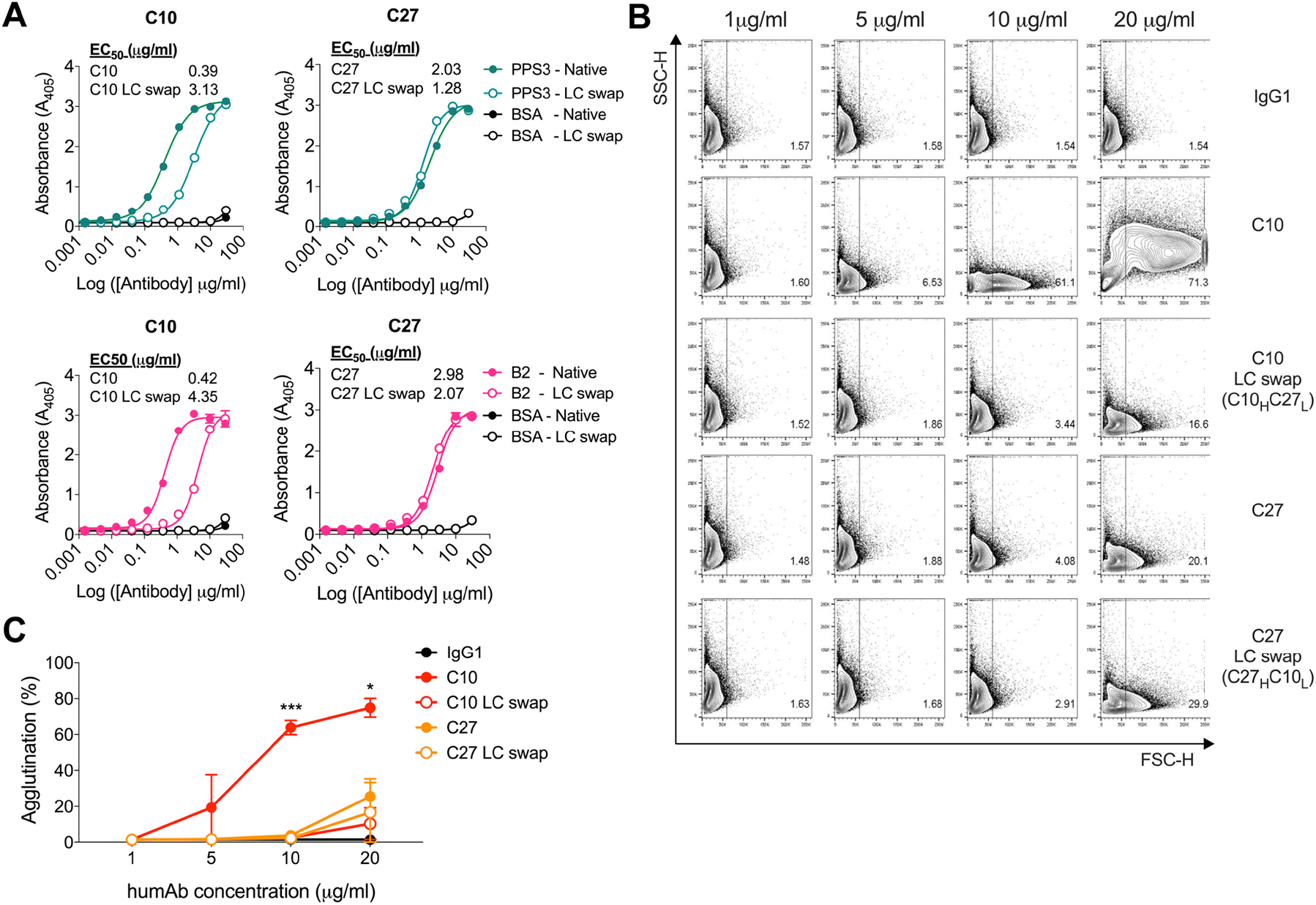
HumAb binding and agglutination of humAbs with light chain swaps. A) HumAbs (native) with their LC swaps were generated and tested by ELISA for binding reactivity to purified PPS3 and B2. Absorbance at 405 is shown on the Y axis for the humAb concentrations shown on the X axis for each humAb. The numerical half-maximal binding titer (EC_50_) is depicted on the graph. Results are representative of 3 independent experiments (n = 2). ST3 strain B2 was incubated with increasing concentrations of humAbs (C10, C10 LC swap (C10_H_C27_L_), C27, C27 LC swap (C27_H_C10_L_)) or control IgG1 and analyzed by flow cytometry. B) Representative FACS dot plots showing percentage agglutination of the indicated native humAb or LC swap at various concentrations. C) Line graph depicting percentage agglutination on the Y axis for concentrations of indicated humAbs and LC swaps on the X axis. Results are representative of 2 independent experiments (n = 2 per condition). By one-way ANOVA; at 10ug/ml; (C10 vs IgG1, C10 vs C10 LC swap (C10_H_C27_L_) C10 vs C27, C10 vs C27 LC swap (C27_H_C10_L_) \*\*\**P*<0.001); at 20ug/ml (C10 vs IgG1, C10 vs C10 LC swap (C10_H_C27_L_), C10 vs C27, C10 vs C27 LC swap (C27_H_C10_L_) \**P*<0.05).

## DISCUSSION

Here we report the gene use and *in vitro* functional activity of seven PPS3 humAbs generated from pneumococcal vaccine recipients. We also demonstrate the efficacy of two humAbs (C10 & C27) which use the same V_H_ and V_L_ genes (V_H_3-9*01/V_L_2-14*03) against NP colonization and lethal ST3 infection in mice. Our data show that the humAbs with the highest affinity, C10, C34, and C38, mediated the most ST3 agglutination and opsonophagocytic activity. Agglutinating PPS antibodies have been reported to enhance complement activation and complement-dependent killing *in vitro* and have also been shown to be important for reducing pneumococcal colonization in mice (20, 22, 23). Notably in our study, humAb ST3 agglutination occurred at low concentrations (≤20 μg/ml), whereas other reported PPS antibodies mediated agglutination of ST14 (100 μg/ml) (22) and ST23 (250 μg/ml) (23) at much higher concentrations. It is possible that humAb agglutination could have augmented CFU reductions in the OPA, as this was observed in the absence of HL60 cells. However, we do not know if this reflected ST3 clumping or killing (26).

Consistent with prior work demonstrating V_H_3 restriction of PPS- and other polysaccharide-binding antibodies (27–30), each humAb except C38 used a V_H_3 gene element. PPS3-specific residues important for PPS23F binding of a V_H_3-30 humAb (31) were not present in our humAbs. However, C10, C27, C38, C11, each had Ala-Arg-Asp: ARD or Ala-Arg-Gly: ARG V_H_ CDR3 motifs, which have been described in PPS-binding (32) and polyreactive antibodies from pneumococcal vaccine recipients (33). There were no common V_L_ motifs, but the C18 V_L_ CDR3 was identical to a PPS8-binding kappa humAb that used the same V_L_ gene (V_L_ 2-30) (32). Serological cross reactivity has not been described for PPS3 and PPS8, but they are similar structurally (34).

In depth analysis of C10 and C27 humAbs revealed that in contrast to C10, C27 had lower PPS3 affinity, minimal agglutinating ability, did not mediate opsonophagocytosis and had more somatic mutations in its V_L_ relative to the germline. Nonetheless, both C10 and C27 reduced NP colonization with ST3 A66 and the clinical ST3 strain, B2 (Table 2). Similarly, both humAbs prolonged survival after lethal A66 IP infection, suggesting that complement and neutrophils in the blood may have enhanced the ability of lower affinity C27 to mediate ST3 clearance, as described for polyclonal IgG (35). However, IN administration of C10, but not C27 was protective against lethal IN challenge with B2. Even though both humAbs reduced NP colonization and inflammatory cytokines in the NP colonization model with this strain, it appears that only C10 prevented dissemination. Notably, an agglutinating mouse mAb, 1E2, prevented dissemination to the lungs after NP colonization, whereas an opsonic mouse mAb, 7A9, did not (13). However, we do not know if the reduced efficacy of C27 in this model reflects an inability to prevent dissemination, and/or distinct features of the ST3 clinical strain, B2. Tissue specific differences in virulence have been identified for other STs (36, 37), but further work is needed to dissect the roles that humAbs and ST3 strain specific differences may play in the reduced efficacy of C27 observed in the lethal IN infection model.

**Table 2.**
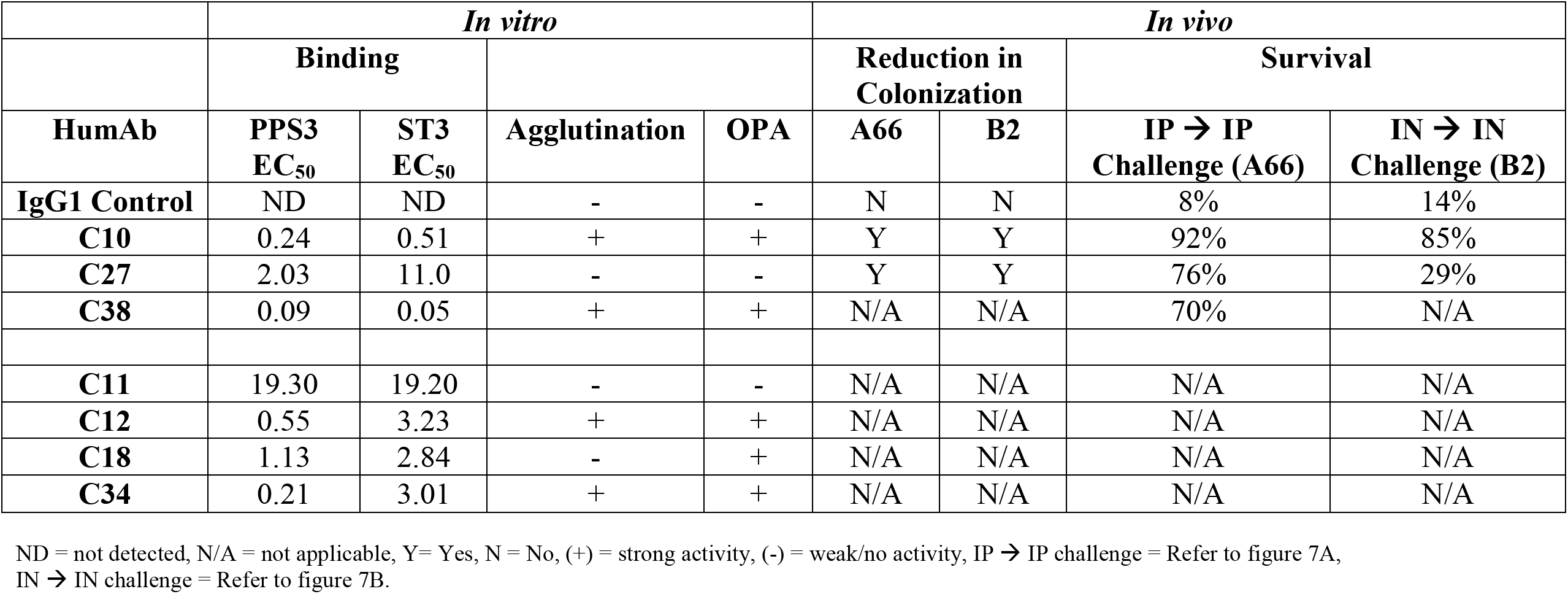
Summary of *in vitro* and *in vivo* functions for all PPS3 humAbs.

The main mechanism by which pneumococcal vaccine-elicited antibodies are thought to confer protection is by mediating ST-specific opsonophagocytosis and this function has been considered a surrogate for vaccine efficacy in clinical studies (6–8). While vaccine effectiveness studies support this association for most vaccine-included STs, this is not the case for ST3 (against which current vaccines are less effective compared to other STs) (2). Given that our data show that C10, which was highly agglutinating and opsonophagocytic, reduced colonization and protected against lethal ST3 infection, its efficacy could stem from its agglutinating ability. There is now ample evidence that ST-specific agglutination can reduce NP colonization in mice (13), but less evidence that opsonophagocytic antibodies reduce colonization. In fact, a PPS mouse mAb (7A9) protected against pneumonia and sepsis but did not reduce colonization in mice (11, 13). Thus, it is possible that ST-specific agglutination, which has not been examined as a correlate of pneumococcal vaccine efficacy in clinical studies, may be a better correlate of vaccine effectiveness against pneumococcal colonization and transmission than opsonophagocytosis. In support of this concept and previously highlighted, agglutinating PPS antibodies are important in prevention of pneumococcal colonization in mice (20, 22, 23). While this may help explain how C10 and C38 worked in our models, it does not explain the efficacy of C27.

Given that C27 did not exhibit agglutination or opsonophagocytosis *in vitro*, but reduced colonization and prevented death from IP infection *in vivo*, it may have mediated these functions *in vivo*. Nevertheless, its lower affinity seems to make this unlikely and we cannot explain its activity based on known mechanisms of PPS antibody action. Thus, we explored the possibility that both C10 and C27 may exert direct effects on ST3 and alter its biological state, as described for a mouse PPS3 mAb that altered gene expression and affected ST3 survival (16, 19). Similarly, we observed a downregulation in *dpr,* which is normally expressed in response to intracellular iron and needed to sequester iron to protect bacteria from oxidative damage (38–40). However, in contrast to the previous *in vivo* study, our *in vitro* data show that C10 and C27 reduced expression of other ST3 genes including *blpX*, an immunity gene needed to avoid bacteriocin-mediated suicide and protect against other bacteriocins (41) and *piuB*, which is essential for regulating iron transport (42). Given their importance in the response to oxidative stress, it is possible that PPS3 antibody-mediated downregulation of these genes could affect ST3 survival. In fact, alteration of ST2 pneumococcal gene expression was reported with penicillin treatment, which similarly reduced expression of genes related to pneumococcal iron uptake (Piu) operon *piuBCDA* and competence (43). Experiments to assess the effect of these humAb-induced changes in ST3 gene expression *in vitro* on ST3 viability *in vivo* are beyond the scope of the current study.

The affinity differences between C10 and C27 could be related to their distinct V_H_ and V_L_ mutations. Notably, for clonally related PPS14 Fabs, the more extensively mutated V_H_ region had lower affinity (44). Similarly, more highly mutated mouse *Cryptococcus neoformans* capsular polysaccharide mAbs had lower affinity and less efficacy *in vivo* (45). Although C10 and C27 have a comparable number of mostly distinct V_H_ mutations, the C10 V_L_ (IGVL2-14*- 03) is closer to germline than C27. Given that the C10 LC swap (C10_H_C27_L_) had lower PPS3 affinity and was less agglutinating than native C10, its superior binding and efficacy against B2 may depend on its V_L_ structure. Notably, structure-function studies of viral antibodies have revealed that V_L_ gene use and structure can dictate whether an antibody is neutralizing or non-neutralizing (46, 47). Our data show that the C10 V_L_ plays a critical role in its agglutinating activity, which was lost when we substituted its V_L_ with the V_L_ of C27. On the other hand, substituting the C27 V_L_ with that of C10 did not alter its agglutinating activity. Together, these findings highlight the potential importance of V_L_ structure and V_H_/V_L_ pairing for PPS3 agglutination, which may depend on a specific PPS3 epitope-humAb interaction. Understanding this interaction will require identification of humAb PPS3 epitopes and structural requirements for binding, as recently reported for a PPS3 mouse mAb V_H_ (21) however with a focus on the V_L_.

To our knowledge, this is the first in depth report of the binding and functional characteristics of pneumococcal vaccine elicited PPS3 humAbs. Our findings reveal an unexpected role for the V_L_ in PPS3 binding and agglutination, and confirm prior reports demonstrating the ability of PPS3 antibodies to affect ST3 gene expression *in vitro*, suggesting a possible mechanism by which non-opsonic and non-agglutinating antibody functions may translate into an ability of certain human PPS3 antibodies to reduce ST3 NP colonization. Although more extensive analyses are needed to understand the impact of PPS3-humAb structure-function relationships on antibody-mediated protection, our data suggest that such investigations will be needed to inform the development of therapeutic ST3 humAbs and more immunogenic ST3 vaccines, which remain urgently needed given the continued threat of ST3 infection globally (1–5).

## MATERIALS & METHODS

### Bacteria

*S. pneumoniae* ST3 strain A66 (provided by David Briles; University of Alabama at Birmingham, AL) and a clinical ST3 strain, B2 (isolated in the Montefiore Medical Center (MMC) clinical microbiology laboratory under Albert Einstein College of Medicine IRB protocol 2014-4035) were grown as previously described (13).

### Mice

6-8 week-old wildtype (WT) female C57BL/6 mice (NCI) were housed in the Albert Einstein College of Medicine Institute for Animal Studies (IAS). All animal studies were approved by the Institutional Animal Care and Use Committee at Albert Einstein College of Medicine (protocol #20171212).

### PBMC blood collection

After obtaining informed consent under Einstein/Montefiore Institutional Review Board protocol 2016-7376, PBMCs were isolated by density gradient centrifugation as described (48) from whole blood of healthy volunteers seven days after pneumococcal vaccination (Pneumovax or Prevnar13). PBMCs were stored in liquid nitrogen prior to use.

### PPS3-PE antigen optimization

Concentrations of fluorescently conjugated PPS3 (PPS3-PE) (Fina BioSolutions) were incubated with ST3 mouse hybridoma cells (11) with or without unlabelled PPS3 (25 µg/well). PPS3-PE positive cells were gated by flow cytometry with cells without PPS3-PE as negative controls. The optimal concentration had similar background fluorescence to control cells (Figure S6).

### Sorting of PPS3-binding memory B cells by flow cytometry

PBMCs were combined from 3 pneumococcal vaccine recipients (two Pneumovax and one PCV13 recipient), to increase probability of isolating PPS3-specific memory B cells. PPS3-memory B cells were defined as (CD19^+^CD27^+^IgM^−^IgG^+^PPS3^+^). PBMCS were stained with PPS3-PE and anti-human fluorescently-conjugated: CD19-PE-Cy7, CD27-APC, IgM-FITC, IgG-V421, CD3-V500, CD4-V500, CD8-V500 and CD14-V500 (BD). Live/dead (LD) cells were identified with Zombie aqua fixable viability kit (Biolegend). CD3, CD4, CD8 and CD14 positive cells were excluded. Gating strategy shown in Figure S7. Single cells were sorted on a BD FACSAria II into 96-well PCR plates (MicroAmp Endura Optical 96-Well Clear Reaction Plates, Life technologies) into lysis buffer as described (49).

### HumAb generation

Variable heavy (V_H_) and light (V_L_) chain immunoglobulin genes from sorted B cells were PCR amplified, sequenced, cloned, and produced as human IgG1s in HEK-293 cells as described (49, 50). For cloning and ligation into human IgG1 expression vectors (IgG-AbVec (PBR322 based), Igκ-AbVec (PBR322 based) and Igλ-AbVec (PBR322 based) (obtained from (50)), refined primers listed in (51) were used to generate DNA fragments with overlapping ends. Gibson assembly was performed to ligate DNA fragments with their corresponding digested vectors using the NEBuilder® HiFi DNA Assembly Master Mix (NEB) according to the manufacturer’s guidelines. Sequencing of V_H_ and V_L_ regions was performed by GENEWIZ (New Jersey, NY). HumAbs were purified using the Gentle Ag/Ab binding and Elution Buffer kit (Thermo Scientific). HumAbs were concentrated using Millipore amicon ultra centrifugal filter tubes (30K MWCO) and resuspended in 200mM NaCl and 20mM Hepes pH 7.4.

### ELISA to determine binding profiles

PPS3 ELISAs were performed using 96-well Nunc Maxisorp plates (Thermofisher Scientific) coated with purified PPS3 (ATCC) (10 µg/ml) in PBS overnight at 4°C as described (11, 52). Pneumococcal polysaccharide 8 (PPS8) (ATCC) (10 µg/ml) was used as a negative control. The numerical half-maximal binding titer (EC_50_) was determined by graphpad prism. A whole-cell ELISA (53) was used to determining binding to the clinical strain B2, similar for PPS3.

### Generation of F(ab’)_2_ fragments

F(ab’)_2_ fragments were generated using IdeZ protease (NEB), purified using CaptureSelect LC-lambda affinity matrix (human) (ThermoFisher), and concentrated with amicon ultra centrifugal filter tubes (30K MWCO) according to manufacturers’ instructions. Digestion and purification were confirmed by SDS-PAGE using mini-PROTEAN TGX pre-cast gels (4-20%) (BioRad).

### *In vitro* agglutination of ST3 bacteria

HumAb agglutination of ST3 was determined by flow cytometry as described (23, 54). ST3 strains A66 or B2 (1x 10^5^ CFU) were incubated with humAbs, F(ab’)_2_ fragments or human IgG1 (control) (Southern Biotech) for 1hr at 37°C in a 96-well plate. Cells were fixed with 1% paraformaldehyde and analysed by flow cytometry. Bacteria were gated on forward (FSC) and sideward (SSC) scatter (referring to cell size and granularity) to determine percentage agglutination. Agglutination was also assessed by light microscopy. Aliquots from each sample were spotted onto 1% agarose pads and visualized with an AxioImager Z1 microscope (Zeiss).

### Immunofluorescence

HumAbs (20 µg/ml) were mixed with 1×10^6^ bacteria (50ul) in microcentrifuge tubes and incubated for 1hr at 37°C. Bacteria were washed 1x with PBS by centrifugation and anti-human IgG-FITC was added to each sample and incubated for 1hr at 37°C. After washing, aliquots were spotted onto 1% agarose pads and visualized with an AxioImager Z1 microscope (Zeiss) (100X magnification).

### Opsonophagocytosis assay (OPA)

The assay was performed with differentiated HL-60 cells at an effector/target cell ratio of 400:1 as described (11, 24). HumAbs and IgG1 (control) (Southern Biotech) were diluted 3-fold from 20 µg/ml. ST3 (A66) killing (%) was determined in the presence of humAbs under 2 conditions: with HL60 cells and complement (3-4 week rabbit complement, Pel-Freez) or without HL60 cells (humAbs and bacteria only), by plating aliquots of samples onto blood agar plates and enumerating CFU.

### *In vitro* bacterial gene expression by reverse transcription-quantitative PCR (RT-qPCR)

To analyse the expression of selected genes during *in vitro* growth as previously described (16), in brief bacteria were grown as described above, diluted to a starting OD of ∼0.01 and 1 ml of culture was incubated with humAbs (C10, C27) or IgG1 control at a concentration of 10 µg/ml for 1.5 hours at 37°C. Bacterial RNA was extracted using the TRIzol Max Bacterial RNA isolation kit (Life technologies) using the manufacturers protocol. RNA was then Dnase treated using the TURBO DNA free kit (Invitrogen) and cDNA was synthesized from 200 ng RNA using the iScript cDNA synthesis kit (BioRad). qPCR was performed using Power SYBR green master mix (Life Technologies) with 10 ng cDNA and 10 µm primers outlined in Table S1 as per manufacturers instructions. Amplification was performed on a StepOne Plus real-time PCR system (Life Technologies) using the following conditions: 95°C for 10 mins, followed by 40 cycles of 95°C for 15 seconds and 60°C for 1 min. Relative expression of genes in humAb treated bacteria was calculated using the threshold cycle (2ΔΔ*CT*) method as described previously (55) using the 16S rRNA gene as an internal control and control IgG1-treated bacteria as the reference.

### Mouse infection experiments

Colonization model: Mice were anesthetized with isofluorane and injected intranasally (IN) with 25 µg of humAbs or anti-human IgG1 (Bxcell) (isotype control) diluted in PBS as described (13). 2hrs after humAb administration, mice were infected IN with either 5×x10^5^ CFU of A66 or 1×10^7^ CFU of B2 in 10ul. CFU were enumerated in the nasal lavage (NL) and lungs at the times specified (24 hrs or 4 days) after infection as described (13). NL cytokines were determined after concentration using the Legendplex Mouse inflammation panel (13-plex) (Biolegend) as per manufacturer’s protocol.

Lethal infection model: Mice were injected either IP or IN with 25 µg humAb or anti-human IgG1 in PBS as described above. 2hr after humAb administration, mice were infected IP with 5×10^5^ CFU A66 (100ul) or IN with 5×10^7^ CFU B2 in 10ul and monitored for survival.

### Nucleotide sequence accession numbers

GenBank accession numbers were as follows: C10V_H_, MZ054262, C11V_H_, MZ054263, C12V_H_, MZ054264, C18V_H_, MZ054265, C27V_H_, MZ054266, C34V_H_, MZ054267, C38V_H_, MZ054268; C10V_L_, MZ054269, C11V_L_, MZ054270, C12V_L_, MZ054271, C18V_L_, MZ054272, C27V_L_, MZ054273, C34V_L_, MZ054274 and C38V_L_ MZ054275.

### Statistical analysis

Data were analysed using a Fisher’s exact test (Survival) or a one-way ANOVA for other analyses as indicated in the figure legends using GraphPad prism. *P*-values ≤0.05 were considered significant.

## ACKNOWLEDGMENTS

We thank Phil Gialanella at Montefiore Medical Center for isolation of the clinical strain B2 used in the study. This study was supported by the National Institutes of Health grants to LP: R01AG045044 and R01AI123654.

## AUTHOR CONTRIBUTIONS

RB designed, performed experiments, analysed, interpreted data and wrote the manuscript. CD assisted with experimental design, contributed to revising and critically reviewing the manuscript. LP supervised the study, designed experiments, interpreted data and wrote the manuscript.

## SUPPLEMENTARY MATERIAL LEGENDS

**Table S1:** Primers used for RT-qPCR

**Figure S1: HumAb binding to ST3 clinical strain B2 by ELISA.**

Binding reflected by absorbance at 405 is shown on the Y axis for the humAb concentrations shown on the X axis for each humAb. Results are representative of 3 independent experiments (n = 2). The numerical half-maximal binding titer (EC_50_) for each humAb is indicated to the right of the panel depicting binding curves of all humAbs.

**Figure S2: Binding of humAbs to the clinical strain B2 by immunofluorescence.**

ST3 clinical strain B2 was incubated with humAbs (20 μg/ml) and antibody binding was detected using IgG conjugated to FITC. Fluorescent images were analysed at 100X magnification and representative of 2 independent experiments (n = 2). Non-specific IgG1 was used as a control. White bar represents 2μM.

**Figure S3: Heavy chain variable region (V_H_) sequences of PPS3-specific humAbs.**

HumAb V_H_ sequences aligned with their germline counterparts based on IMGT/V-QUEST (sequence alignment software). Amino acid changes resulting from somatic mutations are indicated within the sequence alignment.

**Figure S4: Light chain variable region (V_L_) sequences of PPS3-specific humAbs.**

HumAb V_L_ sequences aligned with their germline counterparts based on IMGT/V-QUEST (sequence alignment software). Amino acid changes resulting from somatic mutations are indicated within the sequence alignment.

**Figure S5: *In vitro* agglutination of a clinical ST3 strain.**

HumAbs were tested for their ability to agglutinate a ST3 clinical strain (B2) by flow cytometry. A) Representative FACS dot plots showing the percentage agglutination of all humAbs and control human IgG1 at various concentrations by flow cytometry B) Percentage of agglutination is shown on the Y axis for different humAb concentrations indicated on the X axis. Graph represents data from 2 independent experiments (n = 2 per condition). By one way ANOVA: At 5 µg/ml; (C10, C34 & C38 vs IgG1 \**P*<0.05), at 10 µg/ml; (C10, C12, C34 & C38 vs IgG1 \*\*\**P*<0.001); at 20 µg/ml (C10, C12, C18, C34 & C38 vs IgG1 \*\**P*<0.01), at 40 µg/ml (C10, C12, C18, C27 & C38 vs IgG1 \*\**P*<0.01).

**Figure S6: PPS3-PE optimization with ST3 hybridoma cells.**

ST3 hybridoma cells were optimized with different concentrations of PPS3-PE with and without the presence of unlabelled PPS3 (25 µg/well). PPS3-PE positive signal was determined by flow cytometry. Histograms represent the following groups; Control (cells with no PPS3-PE) (Red), Pre-incubation (cells pre-incubated with unlabelled PPS3 prior to addition of PPS3-PE) (blue), PPS3-PE (cells incubated only with PPS3-PE) (Orange). Results are representative of 2 independent experiments (n = 2).

**Figure S7: Representative gating strategy to sort for PPS3+ Memory B cells.**

PBMCs were collected from patients 7 days following vaccination with pneumococcal vaccines and stained to sort for CD19^+^CD27^+^IgM^−^IgG^+^PPS3^+^ cells (See Methods for details).

